# Allele Sails: launching traits and fates into wild populations with Mendelian DNA sequence modifiers

**DOI:** 10.1101/2024.03.18.585647

**Authors:** Michelle L. Johnson, Bruce A. Hay, Maciej Maselko

**Affiliations:** California Institute of Technology. Division of Biology and Biological Engineering. 1200 East California Boulevard, MC156-29, Pasadena, CA 91125; Applied BioSciences, Macquarie University, North Ryde, NSW 2109, Australia

**Keywords:** gene drive, evolutionary rescue, Cas9, base editing, prime editing, retrotransposon, sex determination, population suppression, population modification, invasive species, malaria

## Abstract

Population-scale genome editing can be used to alter the composition or fate of wild populations. One approach to achieving these aims utilizes a synthetic gene drive element—a multi-gene cassette—to bring about an increase in the frequency of an existing allele. However, the use of gene drives is complicated by the multiple scientific, regulatory, and social issues associated with transgene persistence and gene flow. Alternatives in which transgenes are not driven could potentially avoid some of these issues. Here we propose an approach to population scale gene editing using a system we refer to as an Allele Sail. An Allele Sail consists of a genome editor (the Wind) that introduces DNA sequence edits (the Sail) at one or more sites, resulting in progeny that are viable and fertile. The editor, such as a sequence-specific nuclease, or a prime- or base-editor, is inherited in a Mendelian fashion. Meanwhile, the edits it creates experience an arithmetic, Super-Mendelian increase in frequency. We explore this system using agent-based modeling, and identify contexts in which a single, low frequency release of an editor brings edits to a very high frequency. We also identify conditions in which manipulation of sex determination can be used to bring about population suppression. Current regulatory frameworks often distinguish between transgenics as genetically modified organisms (GMOs), and their edited non-transgenic progeny as non-GMO. In this context an Allele Sail provides a path to alter traits and fates of wild populations in ways that may be considered more acceptable.

## Introduction

The ability to modify or build resilience into ecosystems is increasingly desirable to confront a range of challenges including vector-borne diseases, agricultural loss, the spread of invasive species, and climate change. One important tool to address these are population-scale genetic alterations. These changes can introduce beneficial traits into a population (population modification) or eliminate a harmful population (population suppression). For example, population modification could promote resilience of an endangered or threatened species by bringing currently beneficial genomic modifications to high frequency. It could also be used to introduce ‘anticipatory’ sequence changes designed to provide a benefit in a likely future environment altered by climate change or the introduction of an invasive disease vector. These possibilities are suggested by the growing number of contexts in which one or a modest number of alleles of large effect can produce significant phenotypic benefits for a population. Examples include a single locus that can confer heat resistance in mussels^1^ and cattle^2^, fungi resistance in certain plants^3^, or varroa mite resistance in bees^4,5^. There are also collections of loci that have been suggested to contribute to coral resistance to heat^6,7^, bird resistance to malaria^8^, or frog resistance to chytrid fungi^9^. Genetic alterations can also be used to reduce harm, for example by limiting the ability of insects to vector disease^10,11^, or reducing the toxicity (poison production) of an invasive species such as cane toads. Finally, genetic alterations can be used to bring about population suppression, either directly^12^ or by sensitizing pests to an outside stimulus.

One way to introgress important alleles into a population is via the release of individuals carrying the desired allele. A new allele under positive selection will spread but may take many generations to reach high frequency depending on the size of the benefit and whether it is dominant, additive, or recessive. Alternatively, if the allele is not under positive selection or if the release introduces transgenes meant to suppress the population, large and/or repeated introductions will be needed. These are resource intensive and impractical for many species. Additionally, controlled breeding and the large-scale introduction of these individuals into the target population could increase the frequency of other unintended and possibly harmful alleles. It could also threaten the existing genetic diversity of the target population or subspecies.

These issues can be overcome via the use of synthetic gene-drives (SGD). SGDs are selfish genetic elements that spread to high frequency by biasing their own inheritance; they can include ‘cargo’ in addition to transgenes essential for the SGD’s function. A wide variety of SGDs have been devised (reviewed in (^13,14^)). However, the very features of a gene drive that makes it attractive—that it can rapidly bring to high frequency transgenes that may persist for extended periods in (and in some cases outside) the target region—create regulatory and social hurdles to implementation.

Population-scale genetic alterations with phenotypic consequences can also involve more subtle modifications such as single base changes, or small insertions or deletions. Importantly, in some regulatory environments (Australia provides one example^15^) genome edits present in non-transgenic progeny of a transgenic individual (who bears a DNA sequence modifier such as a Cas9 nuclease) are regulated as non-transgenic. Designation of a population carrying a high frequency of such edits as non-transgenic may facilitate regulatory approval and social acceptance. Here we explore a system that we refer to as an Allele Sail, which can, in the absence of gene drive, bring edits but not transgenes to high frequency for population modification, and cause suppression in certain contexts.

An Allele Sail consists of a chromosomally located genome editor (e.g. a site-specific nuclease such as Cas9, or a base- or prime-editor). We call this editor the Wind, as it is responsible for pushing edits into the population. The editor is expressed in the germline (though the expression need not be germline specific) and introduces sequence modifications at one or more target sites located anywhere in the genome. The homozygous edited individuals are viable and fertile. While the editor is transmitted in a Mendelian fashion, the edits it creates (we call these edits the Sail) increase in frequency at an arithmetic super-Mendelian rate as the editor encounters new unedited alleles in the germline each generation (Figure 1). We use the Allele Sail nomenclature to distinguish its mechanism of action from that of an Allele Pump. An Allele Sail uses a DNA sequence modifier to create *de novo*, *new sequence changes* located at a separate locus. In contrast, an Allele Pump uses a site-specific nuclease and homing^16^ or Toxin-mediated killing of non-carriers of an Antidote in a Toxin-Antidote system^17,18^ to bring about an absolute or relative increase, respectively, in the frequency of *an existing sequence*, usually a transgene located at a separate locus^16^. We first consider the use of an Allele Sail in population modification, where there are a variety of applications for conservation or infectious disease prevention. We then explore potential uses for population suppression in organisms that have sex determination systems in which the activity of a single gene is required for femaleness^19,20^.

**Figure 1.**
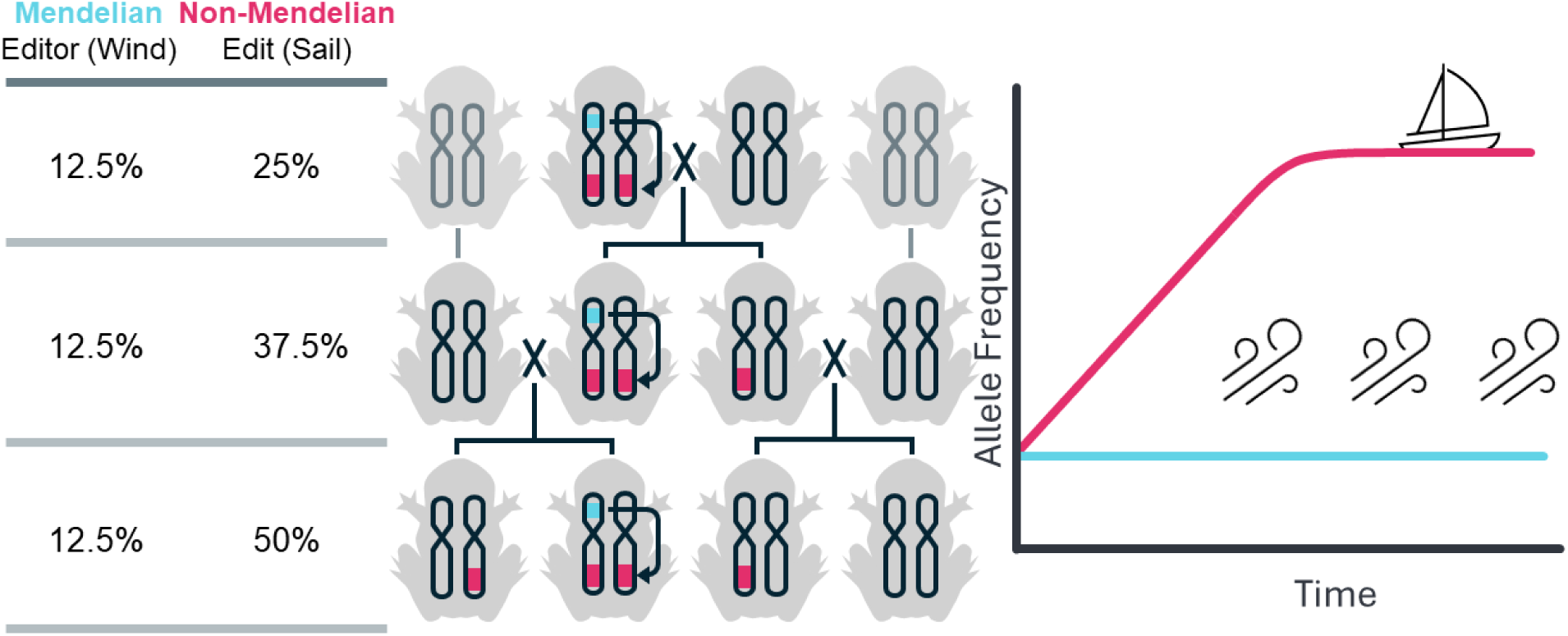
Graphical Representation of Allele Sail Components and Behavior. The Wind (an editor, in blue) is inherited in a Mendelian fashion (left panel). When the editor is present in a germline that carries an unedited target locus located elsewhere in the genome, conversion to the edited state occurs at some frequency (in pink, middle panel). This pushes the Sail (the edited locus, in pink) to higher frequency in the population (left and right panels).

## Results

### Population Modification

We consider the dynamics of an editor that introduces edits at one or more sites in the nuclear genome, resulting in progeny that are viable and fertile. The editor is transmitted in a Mendelian manner, while the edits change in frequency as a function of frequency of the editor, frequency of wild type alleles, fitness costs and editing efficiency. To explore the use of Allele Sails for population modification, we characterize behavior of the components using a discrete-time and generation stochastic model with a panmictic population (see methods for details). This type of model is often used to gain insight into population genetic processes and provides a format that allows comparison of methods for genetically altering populations.

We first consider ideal conditions: the editor alters a target sequence with 100% efficiency, and editing occurs in the male and female germline and in the progeny of a female carrier due to maternal carryover of editing activity. The power of the Allele Sail system can be seen by comparing the frequency of edits over time when introduced in the presence or absence of an editor. These comparisons are shown in Figure 2A for an editor with no fitness cost introduced at a frequency of 10%, with the edit conferring either no fitness change, or an additive benefit or cost of 5% per allele. Recessive and dominant fitness costs/benefits are considered in Supplementary Fig. 1 and show largely similar but distinct dynamics. In the presence of an editor, edits with no cost or a benefit spread rapidly to allele fixation. In contrast, edits conferring a cost rise to high frequency (∼75%) and then decline. In all editing scenarios the peak frequencies of the edits are much higher than those of the corresponding Mendelian allele alone.

**Figure 2.**
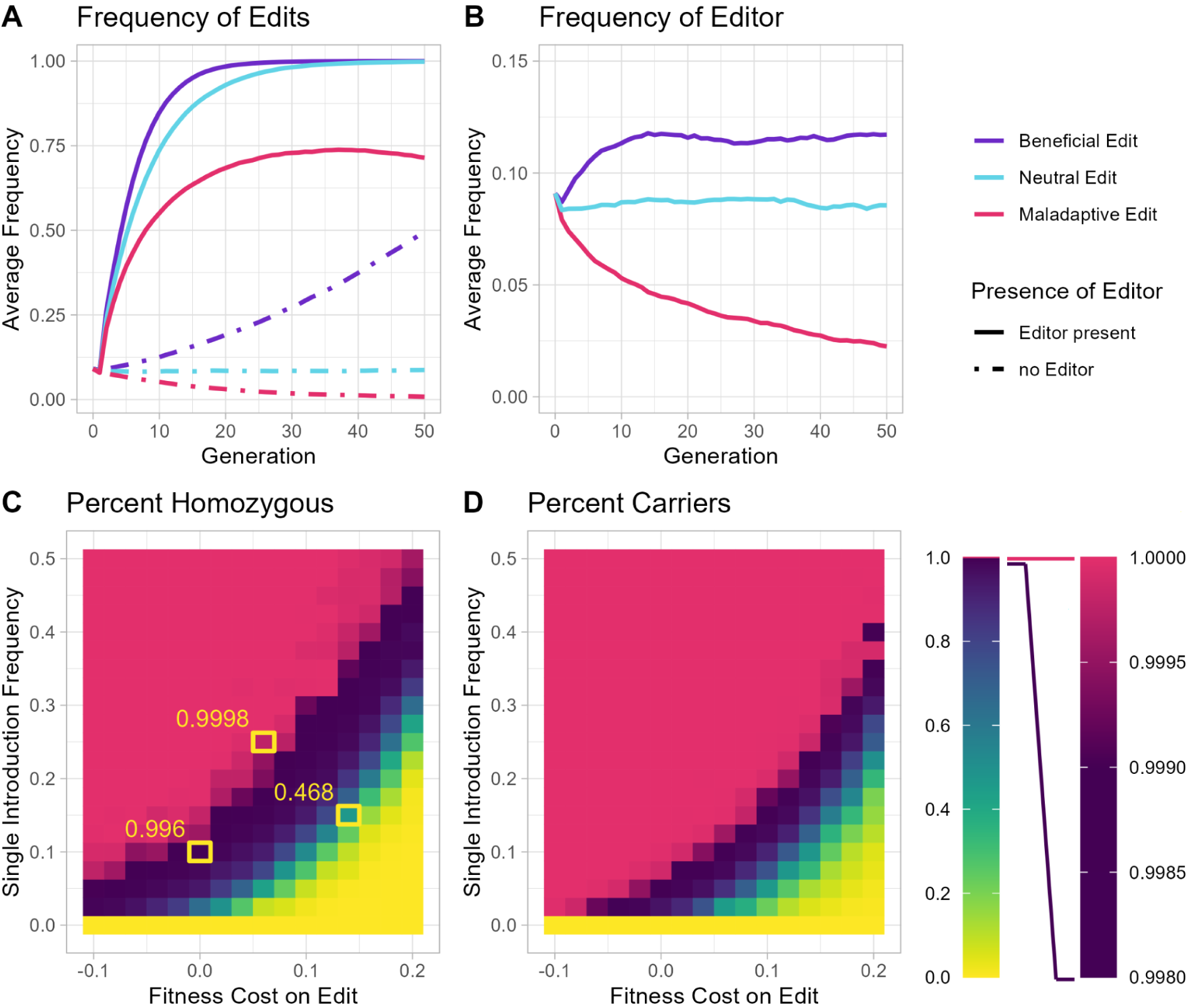
Behavior of a Neutral Editor & Non-Neutral Edit. **A)** The average allele frequency of edits after being introduced at 10% frequency. Introduced individuals are either homozygous for the edit with no editor present, or homozygous for both edit and editor when the editor is present. The fitness benefit shown here is an additional 5% chance of survival for each copy of the edit present (additive benefits), where fitness cost is a 5% decrease in survival for each copy of the edit present. **B)** The average allele frequency of the editor, from the simulations shown in panel **A. C)** Average percent of the population that is homozygous for an edit after 50 generations, for various editor introduction frequencies and fitness costs associated with the edit. Pink tiles represent an average allele frequency of more than 99.9% of the population, across 20 simulations. Negative fitness costs represent fitness benefits, and boxed areas highlight specific allele frequencies to provide guidance for interpretation of the heat map colors. **D)** Average percent of the population that carries at least one copy of the edit, for various editor introduction frequencies and fitness costs associated with the edit.

The dynamics of edit frequency can be understood by considering the fate of the editor (Figure 2B). When the presence of edits has no effect on fitness the editor remains at its introduction frequency, continually generating new edits until fixation is reached. When the presence of the edit confers a benefit the editor increases in frequency. This occurs because the editor spends more time in the presence of the higher fitness edited genotype (which it creates) than does its counterpart wildtype allele. The frequency of the editor plateaus when the edits are ubiquitous (allele fixation) because at this point all individuals have equal fitness. Conversely, when the edit results in a cost to carriers the frequency of the editor declines continuously, since it now spends increased time in lower fitness edited individuals (so long as these never reach fixation) than does its wildtype allele counterpart, leading to its loss through natural selection.

The general relationship between introduction frequency and edit fitness costs/benefits on the frequency of edits is shown in heat maps which plot the frequency of edit homozygotes (Figure 2C) and carriers (homozygotes and heterozygotes) (Figure 2D) at the 50 generation time point. There is a large region of parameter space in which edits are pushed to very high frequency, and increasing the introduction frequency has the general effect of increasing the rate of spread, as well as the time spent at high frequency for those edits that do not reach allele fixation. Finally, we note that the dynamics of edit-bearing genotypes in the population (the frequencies of heterozygotes and homozygotes) depends to some extent on whether editing occurs in the germline with maternal carryover (illustrated above), in the germline only, or in the germline and somatic cells, as well as whether the edit results in a dominant, additive or recessive fitness effects (illustrated for a fitness cost in Supplementary Fig. 2.

The presence of the editor may result in some fitness cost to carriers^21–23^. As such, we explored the context in which the editor but not the edits result in fitness costs. This scenario is illustrated in Supplementary Fig. 3, which shows edit frequency at generation 50 as a function of introduction frequency and additive editor fitness costs, up to 10% per allele. Costs on the editor cause its eventual loss from the population and thus reduce the parameter space in which a single introduction can push edits to high frequency, though increased introduction frequencies and/or multiple releases can compensate. In some cases a guarantee of eventual loss may be desirable, such as when regulatory approval requires that transgenes do not persist in the population. In other contexts, in which the spread of edits to high frequency at minimal cost is the dominant consideration, it may be possible to take advantage of next generation editors that have increased specificity and reduced toxicity^23–26^ to reduce any editor-associated fitness costs.

### Consequences of Altering Editing Efficiency

We have thus-far assumed that our editor has 100% efficiency in editing. CRISPR nucleases such as Cas9 can cleave and create loss of function (LOF) alleles (especially when using multiple gRNAs) in the *Drosophila* germline or the plant *Arabidopsis thaliana* at frequencies near 100%^17,27–34^. While base and prime editors also have significant levels of editing activity in *Drosophila*, they are not 100%; instead closer to 36% for a prime editor^35^, and >90% for a base editor^23^. These encouraging rates notwithstanding, it is important to note that the editing enzymes used—nuclease, reverse transcriptase, deaminase, and uracil-DNA glycosylase—are often derived from organisms that live in temperature ranges very different from those in which their use is intended, which may result in significantly reduced activity in the target species^23,36,37^. To explore these less-than-ideal scenarios, we model several representative examples in which the editor is introduced at a frequency of 10% into the wildtype population and has no associated fitness costs, for various editor efficiencies (between 15-100%), both with and without maternal carryover. These results are compared with a scenario in which edits are introduced directly into the population in the absence of the editor.

Figure 3 shows that decreasing the rate of editing from 100% to 50% or 15% still results in the rapid spread of a beneficial or neutral edit to high frequency by generation 50, as compared with the spread of a Mendelian allele on its own. A deleterious allele also undergoes a significant increase in frequency, though there is ultimately a decline (which also occurs when edits are created 100% of the time) when the frequency of the editor is so low it no longer generates edits faster than they are lost through natural selection. Even so, the peak frequency and time to decline can always be improved by increasing the editor introduction frequency (Supplementary Fig. 4).

**Figure 3.**
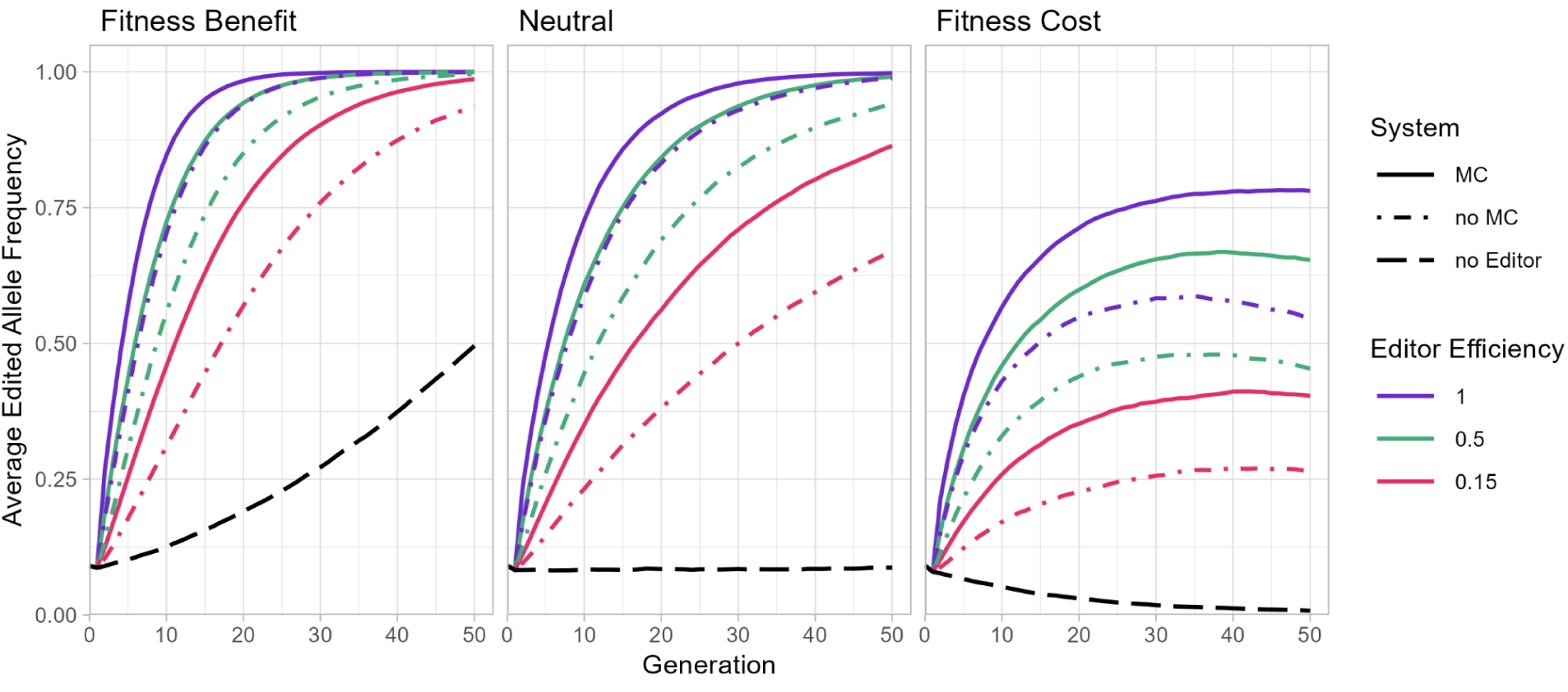
Consequences of reducing editing efficiency for Population Modification. Comparison of edit frequencies over time when using editors with different efficiencies (15%, 50%, and 100%), for various cost/benefits and presence or absence of maternal carryover (MC). Each line represents the average of 20 simulations. Fitness benefits and costs are additive and reflect a 5% increase or decrease in the chance of survival for an individual, respectively. Costs are associated with the edit, and the editor has neutral fitness.

### Consequences of Genetic Linkage Between Editor and Edit

Thus far we have considered scenarios in which the editor and edit site are unlinked. In some cases, particularly if multiple changes are desired, some degree of linkage between the editor and one of the edits may be present. To explore the consequences of linkage we consider the extreme scenarios in which the edit and editor are either tightly linked (are always co-inherited) or unlinked (have equal chance of being co-inherited or not), the editor has a 50% probability of germline editing, and there is no maternal carryover. We utilize a lower rate of cleavage because when editing rates are 100% there is no difference between the linked and unlinked scenarios; all progeny inherit an edit from the carrier parent regardless of linkage. As shown in the heat maps in Fig. 4, for a neutral editor and deleterious edit an absence of linkage is beneficial for spread of the edit (Figure 4). This is because an unlinked editor encounters more non-edited alleles than a linked editor. When fitness costs are associated with the editor, the unlinked version still performs slightly better than the linked (Supplementary Fig. 5).

**Figure 4.**
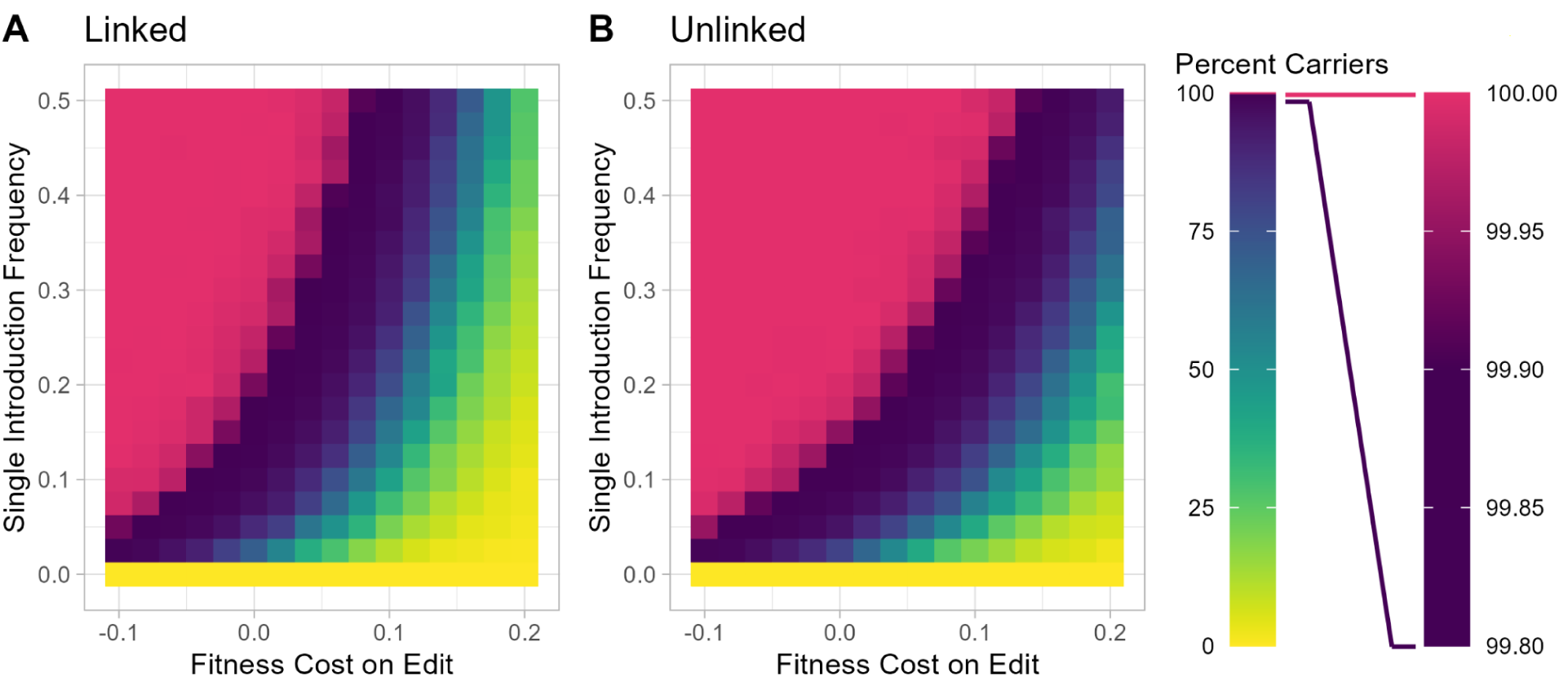
Effects of Linkage between editor and edit on population modification. The frequency of editing is set to 50%, fitness costs are associated only with the edit, and there is no maternal carryover of editing. **A, B)** The percentage of the population that has at least one edited allele at generation 50 averaged over 20 simulations. For higher fitness costs, tight linkage between editor & edit **(A)** results in reduced ability to push edits to high frequency as compared with the case in which editor & edit are unlinked **(B)**.

However, the differences are much smaller and more difficult to see, and beneficial edits spread rapidly regardless of linkage status. In summary, linkage can decrease rates of editing for neutral and deleterious alleles, but the effects are only significant when linkage is tight and editing rates are substantially below 100%.

### Population Suppression

Transgene-based population suppression strategies take several forms. In one, a self-sustaining gene drive utilizes high frequency homing to spread a multi-gene cassette into (thereby inactivating) a haplosufficient gene required in somatic cells for female sexual identity, fertility, or viability. This drives the population towards a homozygous genotype that is fit in males but unfit in females, leading to a population crash^12,38,39^. There are also suppression strategies that do not rely on drive; Non-drive transgene-based approaches utilize periodic inundation of males to, by one mechanism or another, reduce the frequency of female progeny, of progeny generally, or of fertile females^40–44^.

Here we explore how an Allele Sail could be used for population suppression by causing sex ratio distortion. Our focus is on species in which expression of a single gene is needed for femaleness, and whose loss results in conversion of these individuals into fertile males. The Transformer gene in medfly *Ceratitis capitata* provides one example (reviewed in ^45^). Aromatase, encoded by the cyp19a1a gene, plays a similar role in a number of vertebrates. Aromatase converts androgens to estrogens and its loss through chemical inhibition or mutation converts genetic females to fertile males^20,46,47^. This, combined with the fact that estrogen agonists can promote femaleness in genetic males^47^ argues that aromatase activity is both necessary and sufficient for femaleness. Here we model the composition and fate of a population in which an editor such as Cas9 is introduced, creating LOF alleles (the edit) in the aromatase gene. Because single locus sex specification occurs in species with diverse chromosomal systems, we model suppression in species with XY and ZW sex chromosomes. We also remove maternal carryover from these simulations, which allows the editor to persist in the female line for longer. We first consider the consequences of single releases and then multiple releases.

### Dynamics of Alleles and Chromosomes following a Single Release

Results of a single release of the editor at a frequency of 10% carrying capacity are shown in Figure 5A. We compare these to results of a single release of males carrying transgenes that implement female sterile Repressible Inducible Dominant Lethal (fsRIDL), a system in which an autosomal transgene, transmitted in a Mendelian manner, causes death or sterility of female progeny carriers. The suppression effects of fsRIDL are immediate, but the total population also increases back to carrying capacity very rapidly. In contrast, release of an Allele Sail editor results in a relatively prolonged reduction in population size, which is particularly prominent in the ZW system (Figure 5A). To understand why a single release of an Allele Sail with no associated fitness costs only modestly reduced population size rather than collapsing it entirely, we first examined the XY system and the frequency of the editor, the edit, and the Y chromosome over time. The initial release is of XX males homozygous for the editor and edit. This results in a transient spike in population numbers because when these mate with XX females only female progeny are produced, since the editor is germline specific. This increase in females leads to an increase in population, as their progeny include both males and females. Interestingly, the editor undergoes a rapid decrease in frequency, even as the frequency of the edit increases dramatically and the population slowly returns to its carrying capacity (Figure 5B-F). These dynamics (also observed when XY males are released; Figure 5D) can be explained by considering how the frequency of males in the population changes over time. As edits accumulate, males (many now XX) constitute an increasing fraction of the population (Fig. 5A, dashed lines). The editor also spends more time in males than in females as a result of its germline editing activity (Supplementary Fig. 6). Since each female only mates once, males are in excess and the probability that an editor-bearing male will participate in reproduction is reduced. In consequence, the frequency of the editor declines while the edit stabilizes at an intermediate frequency, thereby preventing further population suppression. The Y chromosome (Fig. 5B) is lost from the population for similar reasons. As edits accumulate, XX males make up a larger fraction of the (increased) male population, resulting in a corresponding decrease in the probability that Y-bearing males participate in reproduction.

**Figure 5.**
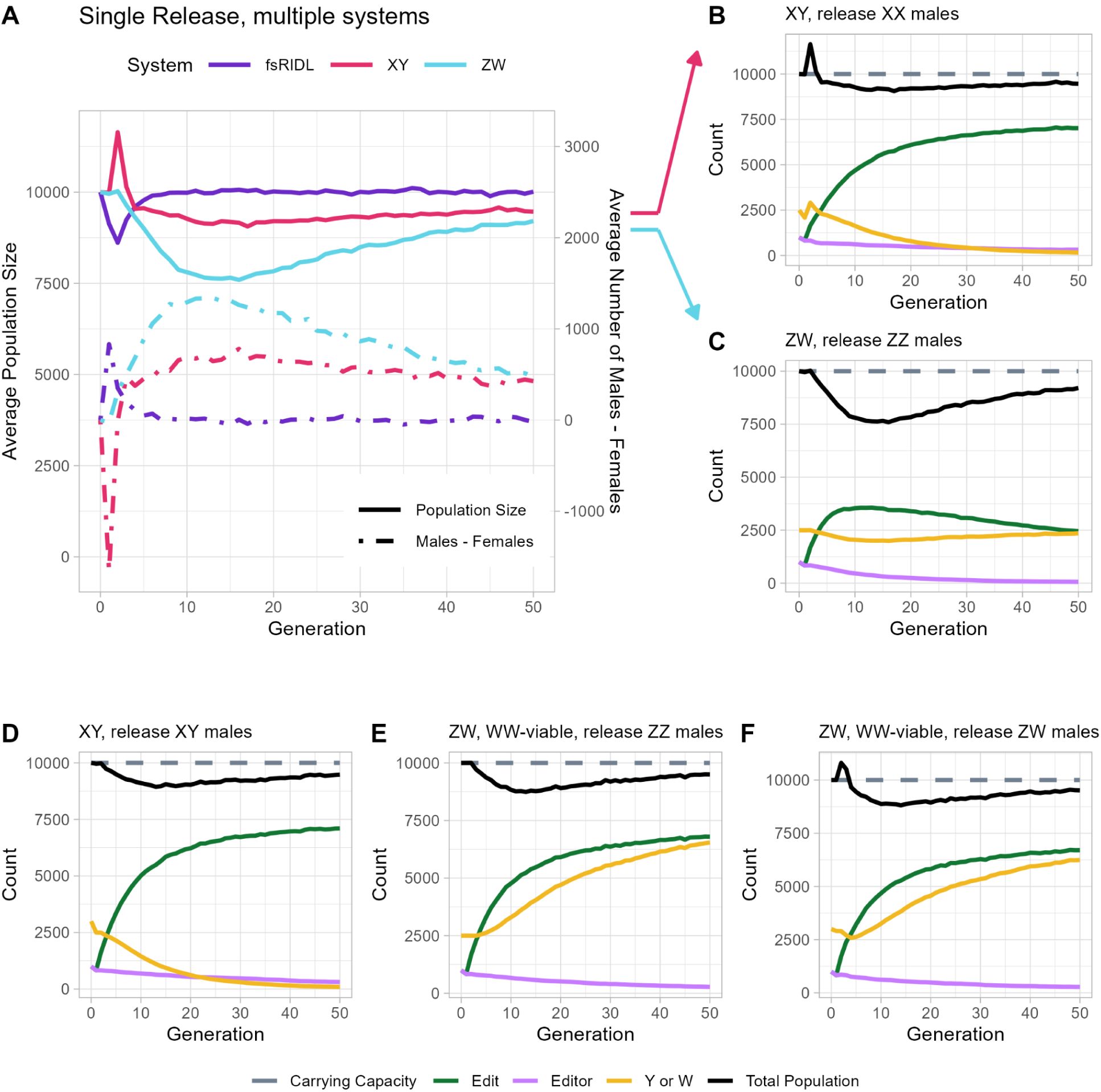
Single Release of Allele Sail for Population Suppression. Values are averaged over 20 stochastic simulations, after a single release of transgenic individuals, with introductions being 10% of the carrying capacity. The counting of population size occurs before this additional release, so those individuals are not counted here. Additionally, there is no maternal carryover occurring in these simulations **A)** An overview of the total population over time, for XY and ZW systems using Allele Sail for suppression, compared to fsRIDL. **B-F)** A breakdown of allele frequency over time, graphed as a proportion of the total population and compared to the carrying capacity. **B)** For an XY system, releasing editor+/editor+, edit+/edit+, XX males. The introduction of additional X chromosomes leads to an initial spike in population, further explored in Supplementary Fig. 8. **C)** For a ZW system, releasing editor+/editor+, edit+/edit+, ZZ males. The increase in Z alleles only increases the effects of our male-skewing editor. WW individuals are considered non-viable, and die off. **D)** Same as B, but releasing XY males, which does not result in a population spike. **E)** The same as C, but WW individuals are viable, leading to an overall increase in W alleles in the population. **F)** Same as E but releasing ZW males instead of ZZ, causing a population spike.

An important consequence of a single modest release of an editor in an XY system is that the Y is completely lost from the population even as the ratio of males to females approaches 1:1 (Figure 5B). This result implies that sex is now determined through a different mechanism. The aromatase loss-of-function edit plateaus at a frequency of 75%, indicating that the aromatase locus is now the primary sex determining locus, with males being edit+/edit+ and females edit+/edit-. This behavior is predicted by earlier modeling of switching of the heterogametic sex through intermediate states in which multiple systems are active^48,49^. It can be understood intuitively by considering the plot in Figure 6, which shows outcomes when populations are seeded with various frequencies of aromatase edits and then followed for 500 generations. Each point represents the simulation endpoint after 500 generations, by which time the sex ratio has approached the Fisherian 1:1^50^. The frequency of the original sex chromosome (either Y for XY, or Z for ZW) is plotted as the X-coordinate, and the frequency of the new sex chromosome, (the edit) is plotted as the Y-coordinate. The trajectory composed of these points represents a path of equilibria by which sex can transition from male heterogametic to female heterogametic without a change in sex ratio^48,49^. For a complete sex determination system turnover, the Y allele must only drop from 25% frequency to 0%, and cleaved aromatase must reach 75%. With sufficient editor, this turnover is easily achieved and sex determination becomes solely dependent on the status of the aromatase locus. The consequences of such transitions during attempts (particularly failed attempts) to suppress a population warrant further study.

**Figure 6.**
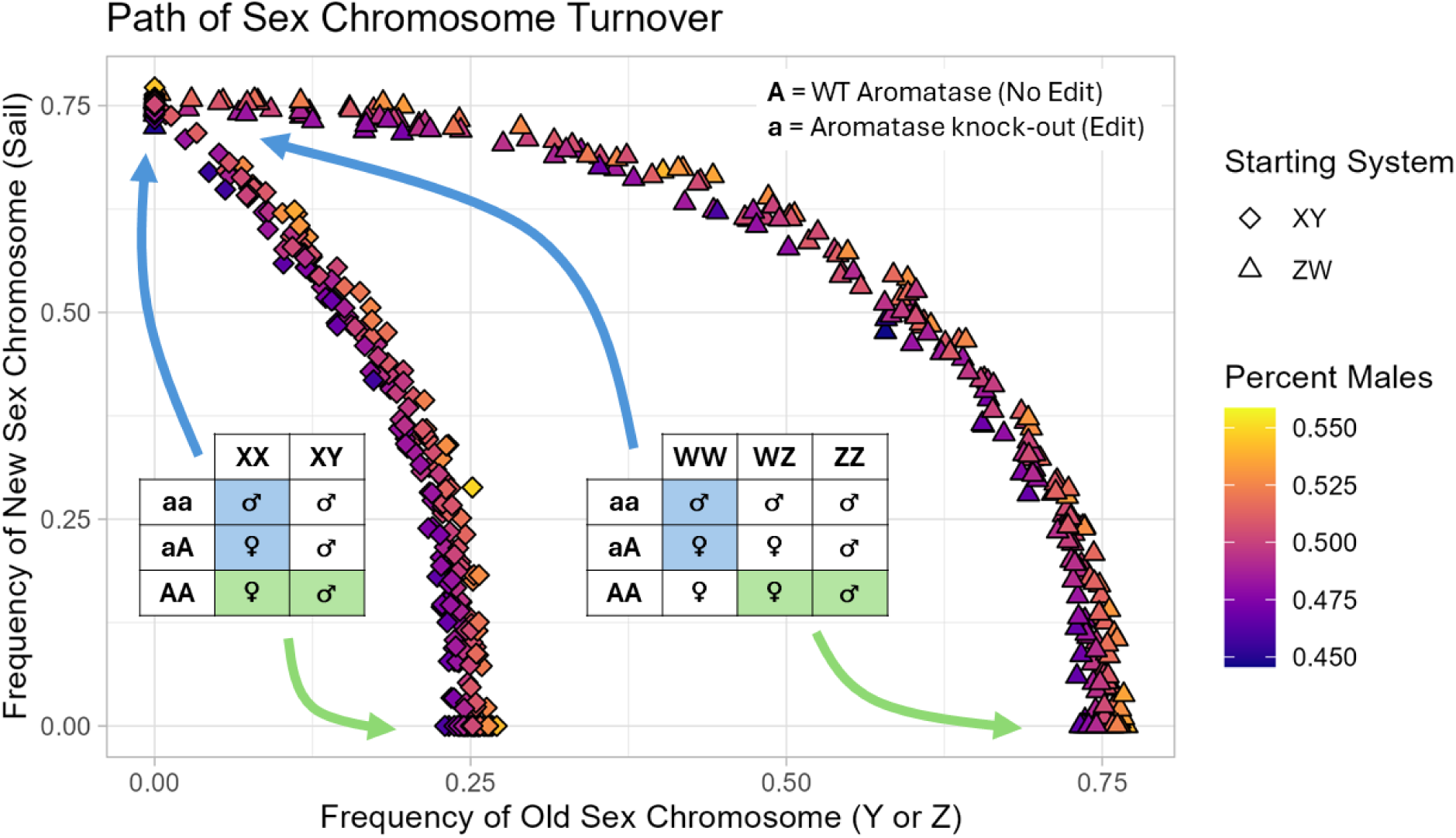
Paths to sex chromosome turnover. Tables represent all possible genotypes within each sex determination system, and their sex. For example, an aaWW individual will be male. A population of the given system, either XY or ZW, was simulated with a single addition of various amounts of Edit (cleaved aromatase). Each point shows the allele frequencies and percent males of each simulation after 500 generations. The frequency of the old sex chromosome (either Y for XY, or Z for ZW) is plotted as the X-coordinate, and the frequency of the new sex chromosome, (the edit) is plotted as the Y-coordinate. WW individuals are viable, allowing for complete turnover from a ZW system in which edits are absent, to a WW system in which the presence or absence of the edit determines sex. The genotypes of these endpoints are shown in the green and blue boxes. The arcs defined by all points represent continuous paths of equilibria connecting two different sex determining systems in which sex ratios remain near 1:1 males:females.

We now consider a ZW system in which the W chromosome is required but not sufficient for femaleness/female viability (e.g. certain birds^51^, crustaceans^52^, and amphibians^53,54^) (Figure 5A, C). Release of ZZ males brings about a gradual but transient decrease in population size (Figure 5A, C), coupled with a transient increase in the male:female ratio (Figure 5A). These effects are due to a rise and subsequent fall in the frequency of edits (Figure 5C). The editor is ultimately lost for the same reasons as in the XY system: it is present more often in males than females (Supplementary Fig. 6), reducing its likelihood of participating in mating. When the WW genotype is inviable, the edits also ultimately decrease in frequency because they find themselves in inviable WW individuals more often than do the non-edited alleles. Placing the editor on the Z-chromosome can mitigate this effect, leading to increased suppression (Supplementary Fig. 7). In the XY system there is no equivalent lethal genotype since the creation of XX males drives the population towards an all XX genotype in which sex is now determined by the presence or absence of a functional aromatase allele. Also unlike the XY system, the W sex chromosome (in contrast to the Y) is not lost from the population because it is required (but not sufficient) for femaleness. Because of this, and because the Z is required for viability, neither can be lost and heterogametic sex chromosome flipping cannot occur. In such a system the editor is quickly lost, edits decrease over time, and the W allele returns to a frequency of 25% (Figure 5C).

In the case where WW homozygotes are viable^52,54,55^ (as might be the case with a newly derived sex chromosome) heterogametic sex flipping can occur^49^, but is harder to achieve than in the XY case. The dynamics that support this conclusion are shown in Figure 5E, F and Figure 6, focusing on the behavior of the W chromosome. Figure 5E and Figure 5F show the consequences of a single release of ZZ or ZW individuals, respectively. In both cases the transient drop in population size is coupled with a substantial and persistent rise in the frequency of the edits and the W chromosome. Both plateau as the editor is lost from the population through the mechanisms discussed above.

The reason the population returns to its carrying capacity even while edits remain at high frequency is due to the fact that WW individuals are viable and can be either male or female depending on the status of the aromatase gene. This point is illustrated in Figure 6, which shows that there is a large region of parameter space in which ZZ males and ZW females coexist at a near 1:1 sex ratio with ZW males and WW females. The presence of edits drives the population towards a WW state (with aromatase status again determining sex), but a much higher frequency of edits is required than with the XY system, since the Z allele must decrease in frequency from 75% to 0% (as opposed to a drop of the Y from 25% to 0%) for the transition to be complete. Thus, a modest single release is insufficient and leaves the population in a state with mixed sex determination systems. These results correspond to previous findings that modeled a scenario termed mildly male-determining XY to ZW turnover^48,56^.

### Dynamics of Alleles and Chromosomes following Multiple Releases

Here we model the effects of repeated releases - once every generation. As illustrated in Figure 7, repeated releases of an aromatase editor at a frequency of 10% eventually causes population collapse (Figure 7A). This contrasts with repeated releases of fsRIDL, which at the same introduction frequencies only decrease the total population size (Figure 7A). To explore multiple release scenarios in more depth we investigated the time to collapse. Plots of the average generation to collapse versus introduction frequency are shown in Figure 7B. An editor is much more effective than fsRIDL for low-frequency releases, causing collapse for releases between 10% and 20%, while fsRIDL only results in collapse when the release frequency is 25%, and even then, only after roughly an additional 10 generations. For higher introduction frequencies, the time to collapse is comparable (Figure 7B). Reduced editing efficiencies can still lead to population collapse under some scenarios. For example, an XY system can still be collapsed faster than fsRIDL (assuming 100% female killing with fsRIDL) when the editing efficiency is 90%, but not when the editing efficiency is 80%. Meanwhile, in a ZW system in which WW individuals are non-viable, an editing efficiency of 80% still promotes population collapse faster than with fsRIDL. (Supplementary Fig. 9).

**Figure 7.**
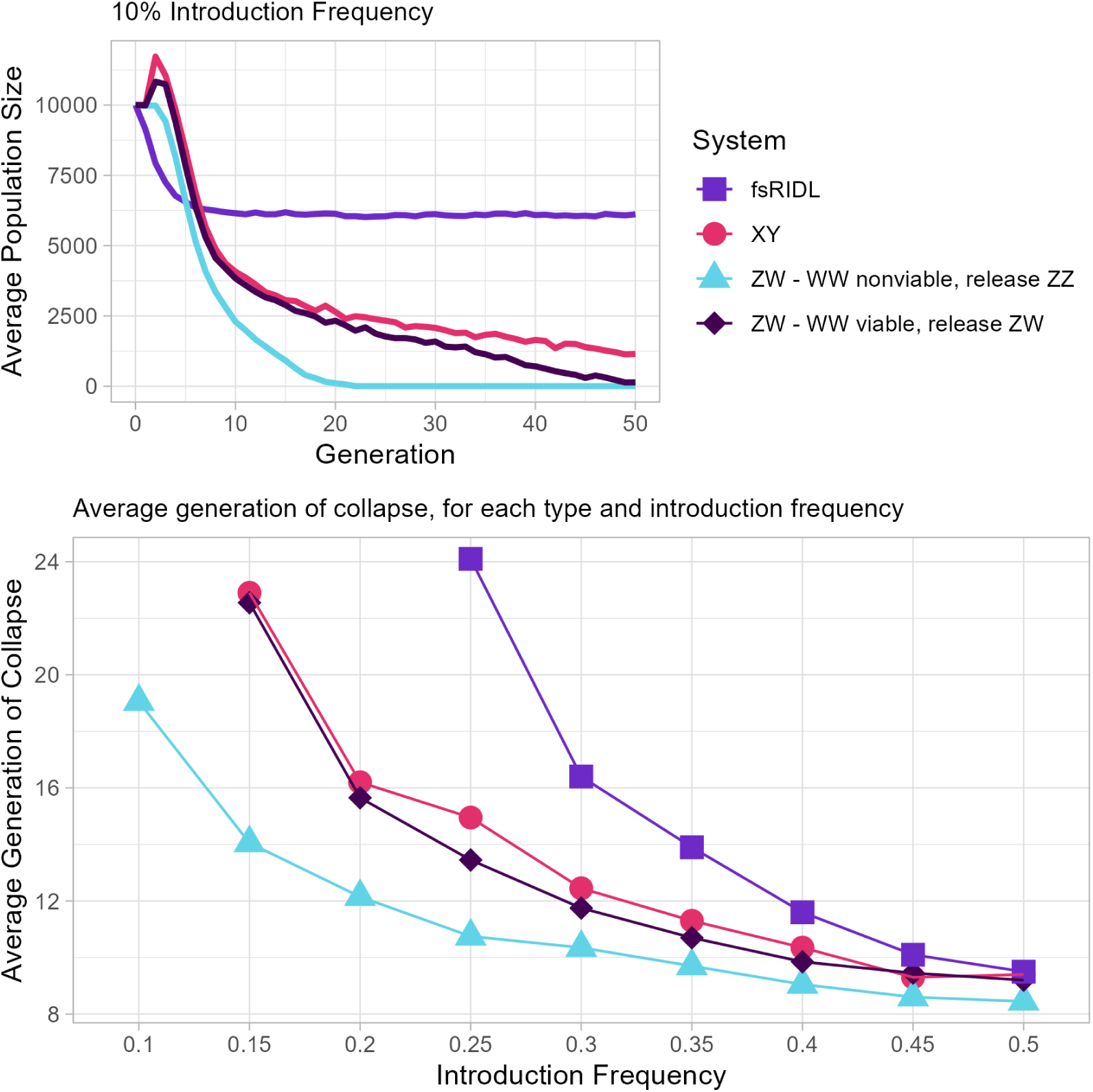
Comparison of population suppression using multiple releases of an Allele Sail or fsRIDL. **A)** Total population over time, averaged over 20 simulations. Transgenic individuals are introduced at 10% of carrying capacity, at the start of every generation. The total population count shown here does not include this additional population, only counting the number of surviving offspring. The XY system shown here introduces XX males, and the ZW introduces ZZ, with WW offspring being either non-viable or viable. **B)** The average generation of collapse for various introduction frequencies. Points plotted here are the average of 20 simulations, where all 20 simulations went to collapse within 50 generations. If all 20 simulations did not collapse within 50 generations, the corresponding point is not plotted. Transgenic individuals were introduced at the indicated introduction frequency, once every generation, until collapse. For introduction frequencies below 25%, an editor can collapse a population where fsRIDL cannot. At higher frequencies, the ability to collapse a population and time to collapse are comparable for strategies using fsRIDL or an editor.

Finally, we asked what the effects on population suppression are if the editor, in addition to cleaving a gene required for femaleness, also creates LOF alleles in a haplosufficient gene required in somatic cells for female viability or fertility - a strategy sometimes referred to as sterilizing sex conversion^57^. As illustrated in Supplementary Fig. 10, sterilizing sex conversion leads to faster collapse, bringing about elimination of a population using repeated releases of only 5% of the starting population (Supplementary Fig. 10).

## Discussion

We set out to investigate the potential of a novel population-scale approach to genome alteration, which we call an Allele Sail. An Allele Sail consists of a genome sequence modifier (the Wind) transmitted in a Mendelian manner, that introduces edits into a target locus (the Sail). A single release of an editor can push a target nuclear allele to very high frequency, even in the presence of fitness costs, on either the editor or the edit. These observations suggest an Allele Sail can be a promising tool for population modification. We note that editors could also be used to alter the genomes of mitochondria and chloroplasts^58^, so as to tune interactions between components encoded by nuclear and organelle genomes that determine fitness^59^. Here we focused on nuclear genome editing. The Allele Sail is a relatively simple system that is applicable for many species. It will also often be self-limiting when the presence of the editor or edits result in costs to carriers. Even in populations where a beneficial allele is already present at low frequencies, an Allele Sail can push that trait into the population much faster than natural selection alone, depending on how many transgenic individuals are released and the efficiency of the Wind/editor. One concern may be that simple editors can only make small changes to the genome. However, small variations such as point mutations or small indels at one or a modest number of loci can have a large effect. As examples, point mutations have been found that contribute to plant disease resistance^3^, animal heat tolerance^1,2^, and honeybee sensitivity to Varroa mites^4,5^. As another point of reference, as of 2019, half of the pathogenic variants in the human ClinVar database were point-mutations, and almost 90% of clinically relevant insertions and deletions were less than 30bp^60^. Finally, recent work shows that larger fragments of DNA (which would albeit be considered transgenes) can be copy-pasted from one location to another using engineered retrotransposons^61,62^. These observations suggest it may soon be possible to push larger fragments of DNA into a population in a self-limiting manner using an editing locus transmitted in a Mendelian manner

### Use Case - when beneficial alleles already exist

One exciting application of an Allele Sail is evolutionary rescue, the process by which introduction of new alleles into a threatened population allows it to adapt rapidly enough to survive a current or anticipated stress. In species where beneficial alleles exist in some populations but not others, two common methods of introducing favorable alleles into the threatened population are selective breeding and targeted gene flow. Examples include breeding toad-smart quolls^63^, and introducing heat-tolerant corals into sensitive populations^64^. However, selective breeding is time and labor intensive, and can reduce genetic variation. Translocation of individuals can also lead to outbreeding depression, while the rapid spread of the beneficial trait is not guaranteed. Genomic editing utilizing an Allele Sail may prove useful because the introduction frequencies can be low, bringing about an increase in trait frequency without dramatically modifying the genetics of the population at other loci.

### Use Case - when beneficial alleles *don’t* already exist

In populations in which the desired beneficial allele is not present, and a future threat is clear, an Allele Sail could be used for anticipatory conservation: pushing into a population an allele that is not currently strongly beneficial but that is projected to be so in the future. In these scenarios, simply introducing or re-introducing a beneficial allele into a threatened population may not be enough to prevent population collapse. If the population drops below its stochastic threshold before the allele reaches high frequencies, the presence of a beneficial allele at a low frequency may not save the population from extinction, or allow the population to persist in the face of rapidly changing conditions^65^. An Allele Sail can dramatically increase the spread of desired alleles before populations begin to experience serious pressure, slowing or preventing population decline to critical levels. An example of this challenge is highlighted by research into avian malaria in Hawaii; small releases of disease-resistant individuals into a population would likely face genetic dilution if the disease pressures are not yet severe, which might prevent the creation of a large, robustly disease-resistant population^66^. Increased transmission of resistance via an Allele Sail could combat dilution and greatly increase the likelihood of success.

### Use Case - non-rescue

An Allele Sail can also spread alleles that are not beneficial to carriers but are beneficial to others. For example, in an invasive species such as Cane Toads, an Allele Sail could reduce the negative effects of the spreading population; if Cane Toads were engineered to produce less toxin or be completely non-toxic, their invasion would be much more survivable for local fauna. An Allele Sail could also spread an anticipatory maladaptation, seeding an increased susceptibility to certain toxicants before introducing said toxicants into a population, thereby increasing its effectiveness. An Allele Sail in which edits result in an immediate fitness cost will select for resistant (WT in phenotypic effect but unmodifiable) sequence variants, as with gene drive for population suppression (reviewed in ^13,14^). In cases where a base or prime editor is used to generate small indels or substitutions it will be important to understand the frequency with which such resistant alleles might arise, either due to bystander sequence editing activity or existing sequence polymorphisms) and their consequences for the intended population effect. In other cases, in which creating LOF alleles is the goal, available evidence suggests it may be possible to prevent resistant allele appearance through targeting of multiple sites in the target gene(s) using gRNA multiplexing^17,27–34^.

### Suppression

In the context of population suppression, we focused on using repeated introductions of an Allele Sail to bring about sex ratio distortion by creating LOF alleles of a haplosufficient gene required in somatic cells for femaleness. In this context an Allele Sail is (when modification rates are high) generally more effective than the non-drive fsRIDL introduced under similar conditions. One important finding is that Allele Sails are self-limiting, even when their presence does not result in a fitness cost, as they remove themselves from the population over time. Even if small numbers of Allele Sail individuals spread through migration into a secondary population, they would not seriously impact total population size. A chemical sex-distortion scheme, with some similarities to the one discussed here, is being tested for Brook trout in Idaho. YY males are created through crosses that utilize XY males and XY females generated by estrogen supplementation. Release of resultant YY males results in all progeny being XY males.^67^ Modeling suggests that regular releases, when combined with yearly 50% suppression via conventional means, have an 80% chance of eradicating brook trout within 9 years^68^. Using an Allele Sail could drastically reduce the number of fish that would need to be released, and decrease the eradication time. Finally, if the editor creates a second set of edits that induce female sterility, the power of the system to bring about population elimination at low introduction frequencies is enhanced. An added benefit of this strategy is that sterilizing sex conversion likely has a higher tolerance to evolution of resistance than non-sterilizing sex conversion^57^.

### Altering Sex Determination

In our investigation of population suppression we encountered an interesting result concerning sex determination. By introducing a modest frequency of an Allele Sail that cleaves aromatase and creates fertile XX males in an XX female; XY male species, we were able to switch a population from having heterogametic males to heterogametic females, creating a new sex determination system. The sail we introduced, cleaved aromatase, is considered a mildly male determining mutant, matching up with Table 2B in Bull & Charnov^48^. The sex determination system turnover happens rapidly in our simulations, as the Y allele is nearly eliminated by generation 50. Introduction of the same frequency of editors into a ZW system did not yield a switch, but could (when WW is viable) do so under multiple release scenarios in which population elimination failed. The ZW to edit+/edit- switch is a much slower process, highlighted by the neutral equilibrium path lengths of these two systems.

### Summary/Conclusions

The differences in behavior of different sex determination systems means that special consideration should be taken for different species. A system where WW individuals are non-viable could be highly efficient, which could be the case in many bird species due to degeneration of the W chromosome^51,69^. Birds are a common agricultural pest, and many attempts have been made to reduce or eliminate invasive birds. Examples include Monk Parakeets and Sacred Ibis in the US^70^, and the 20 exotic species of birds currently in Australia which are known to be agricultural pests^71^. In contrast, there are many species for whom WW individuals are viable, including crustaceans^52^, amphibians^53,54^, and reptiles^55^. When designing for these systems, sterilizing sex conversion may be necessary to achieve the desired outcomes. In cases where an XY system is involved, sex determination system turnover is a legitimate concern and the impacts of turnover would need to be seriously considered.

Despite these limitations, Allele Sails could be useful for suppression across many species. Unlike homing-based drives, the editors in Allele Sail systems, which would only need to create LOF mutations, would not need to rely on homology directed repair mechanisms, and therefore may have a much broader range of use since resistant allele formation can be limited through gRNA multiplexing. When used for suppression, Allele sails are also non-lethal, addressing concerns of humaneness^72^. An additional benefit of Allele Sails is that, in theory, the number of organisms bearing Cas9 or some other editor should never significantly increase in the population. In countries like Australia, offspring that only inherit an edit, but not an editor transgene, could be considered non-GMO, reducing regulatory hurdles^15^. Because of its broad applicability, both in terms of population suppression and modification, we believe Allele Sails are a potentially useful tool for population control, both in reducing and eliminating invasive or harmful species, as well as saving or conserving endangered native species.

## Methods

All modeling was done using a stochastic discrete-generation simulation, for populations with carrying capacity of 10,000 individuals, and individuals that are expected to bear 100 offspring each. The actual number of offspring is pulled from a poisson distribution.

Introduction frequencies represent a release of homozygous males, equal to that percent of the carrying capacity; they are an additional release of individuals on top of the carrying capacity. For example, a 10% introduction frequency of transgenic individuals will have a starting breeding population of 10,000 wildtype individuals, and 1000 transgenic individuals for a total population of 11,000. However, our measure of Total Population, as shown in the figures, does not consider the individuals artificially added to the population and does not count the added individuals when performing population-dependent growth.

Density-dependent growth was implemented as a survival modifier on all offspring; when the population size is equal to carrying capacity, each individual in a litter of size N has a 2/N chance of survival. This should lead to perfect population replacement when there are no fitness costs, as each mating produces 2 offspring. We modify this value to create density dependent growth following the Beverton-Holt model, a discrete-time version of the logistic growth curve. As such, we modify the 2/N chance of survival by a density-dependence factor (eq. 1), where *P* is the current population size, *K* is the carrying capacity, and *g* is some arbitrary growth factor. We used *g = 10*, which means that if population size is near zero, each offspring has roughly a 20/N chance of surviving.

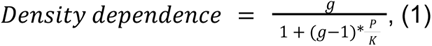

Fitness costs further modify the 2/N chance of survival, such that a 5% fitness cost leads to an individual having a 2/N * 0.95 chance of survival, and a 5% fitness benefit leads to an individual having a 2/N * 1.05 chance of survival. This approach to both fitness and density dependence is based on similar modeling work^73^. Of note, this simulation implements density-dependence in the adult stage. In insects, population size tends to be more dependent on larval density as opposed to adult population density^74^. For these species, our model likely predicts suppressive techniques to be more effective than they are in real life, as cage trials using fsRIDL do not use 1:5 transgenic:wildtype releases, but instead closer to 7:1 transgenic:wildtype^75^. However, we believe that fsRIDL and Allele Sail should be equally affected by this choice, and therefore comparisons between them should hold.

Some simulations included maternal carryover modification and some did not - generally, all modification simulations used maternal carryover of the editor except where noted otherwise, while suppression simulations did not. These differences are noted in the text and figure legends. Maternal carryover, when present, was assumed to be 100%, meaning that if the mother carried an editor, both target alleles in the offspring would be edited regardless of whether the father carried an editor.

The simulation and more information about the code and how to use it can be found at https://github.com/HayLab/AlleleSail.

## Data Availability

All data is available in the main text, the supplementary information files, or in the github: https://github.com/HayLab/AlleleSail

## Acknowledgements

Special thanks to Tobin Ivy for laying the groundwork for this simulation.

## Contributions

Conceptualization: M.L.J, B.A.H. and M.M. Experimental design: M.L.J, B.A.H. and M.M. Coding, modeling, and figure creation: M.L.J. Writing: M.L.J, B.A.H. and M.M.

## Corresponding authors

Correspondence to

## Ethics declaration

The authors declare no conflicts of interest.

## Extended Data

**Supplementary Figure 1.**
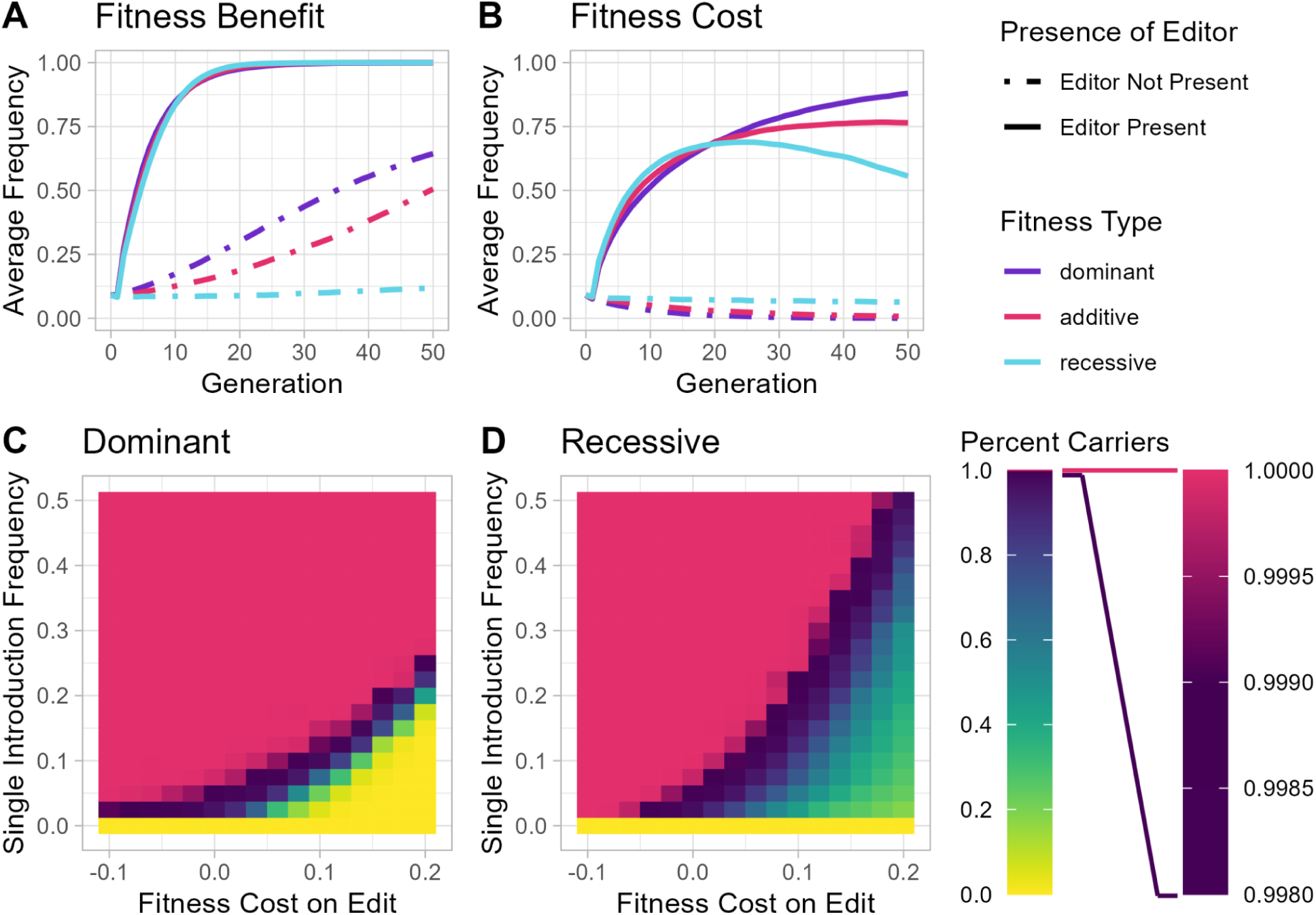
Behavior of Allele Sail edits in response to different types of fitness costs: dominant, additive or recessive. Fitness costs and benefits are associated with the edit. For the dominant and recessive types, fitness costs/benefits are 10% (either positive or negative), and in the additive case are ‘5%’, such that a heterozygous individual has a cost/benefit of 5%, and a homozygous individual has a cost/benefit of 10%. **A)** The average frequency of the edit, both with and without an editor, when transgenic (bearing the edit, or or the edit and editor) individuals are introduced at 10% of carrying capacity and the edit has a 10% (dominant/recessive) fitness benefit. **B)** The same as A, but for a 10% fitness cost. **C)** A heatmap showing the relationship between introduction frequency and fitness costs, when the cost/benefits are dominant and apply to the edit. The percent carriers being plotted here is taken from Generation 50. **D)** The same as C, but for recessive cost/benefits.

**Supplementary Figure 2.**
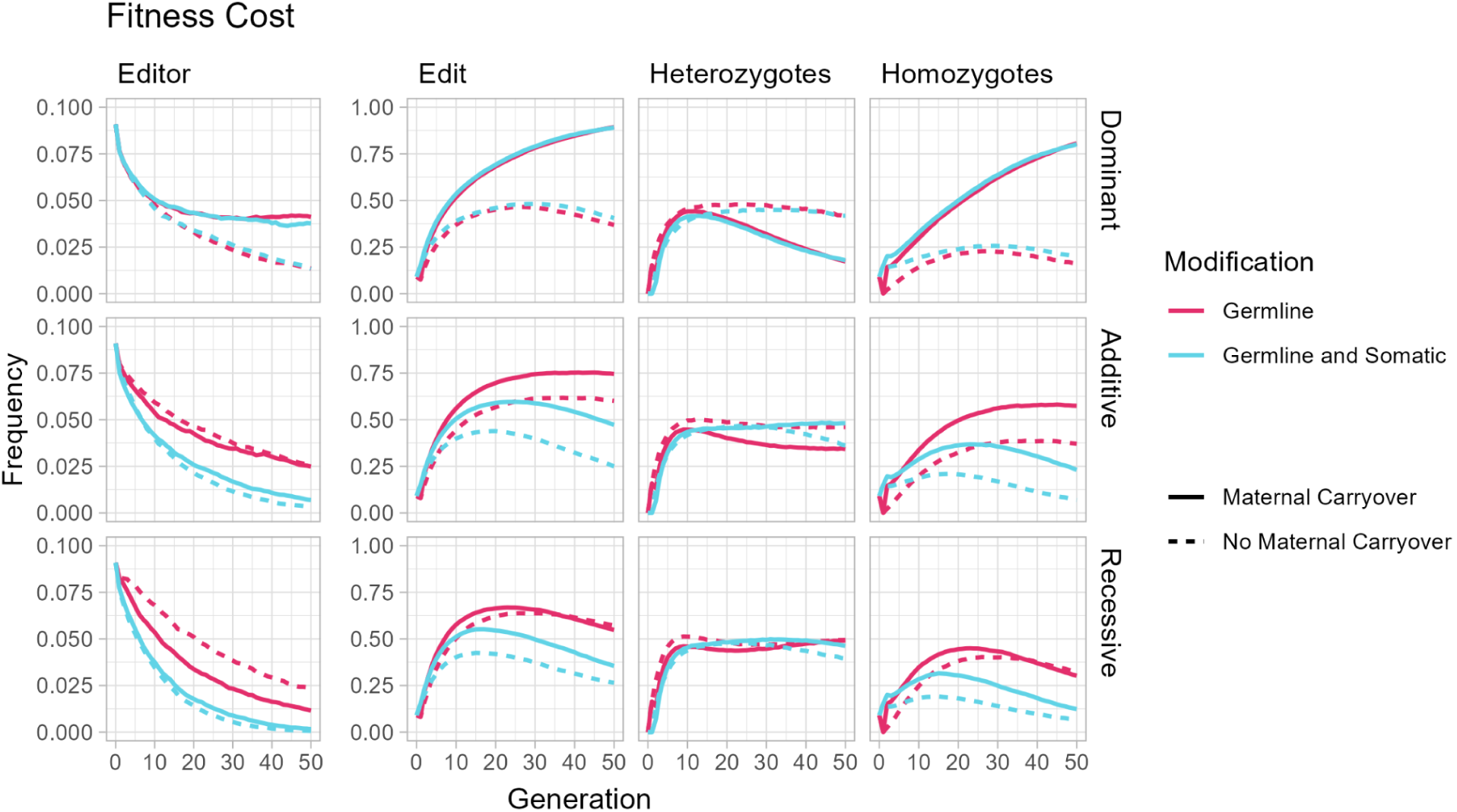
Behavior of Allele Sail edits in response to different types of fitness costs and different modification times. The allele frequency of edits and the editor averaged over 20 runs, along with the frequency of heterozygotes and homozygotes for the edit. This is for different fitness types (Dominant, Additive, and Recessive) as well as for different modifications, either Germline or Germline with Somatic, and also with and without Maternal Carryover. Maternal Carryover refers to Maternal Carryover of the editor, which leads to somatic editing in all offspring of an editor-bearing female. This includes offspring that do not carry the editor themselves.

**Supplementary Figure 3.**
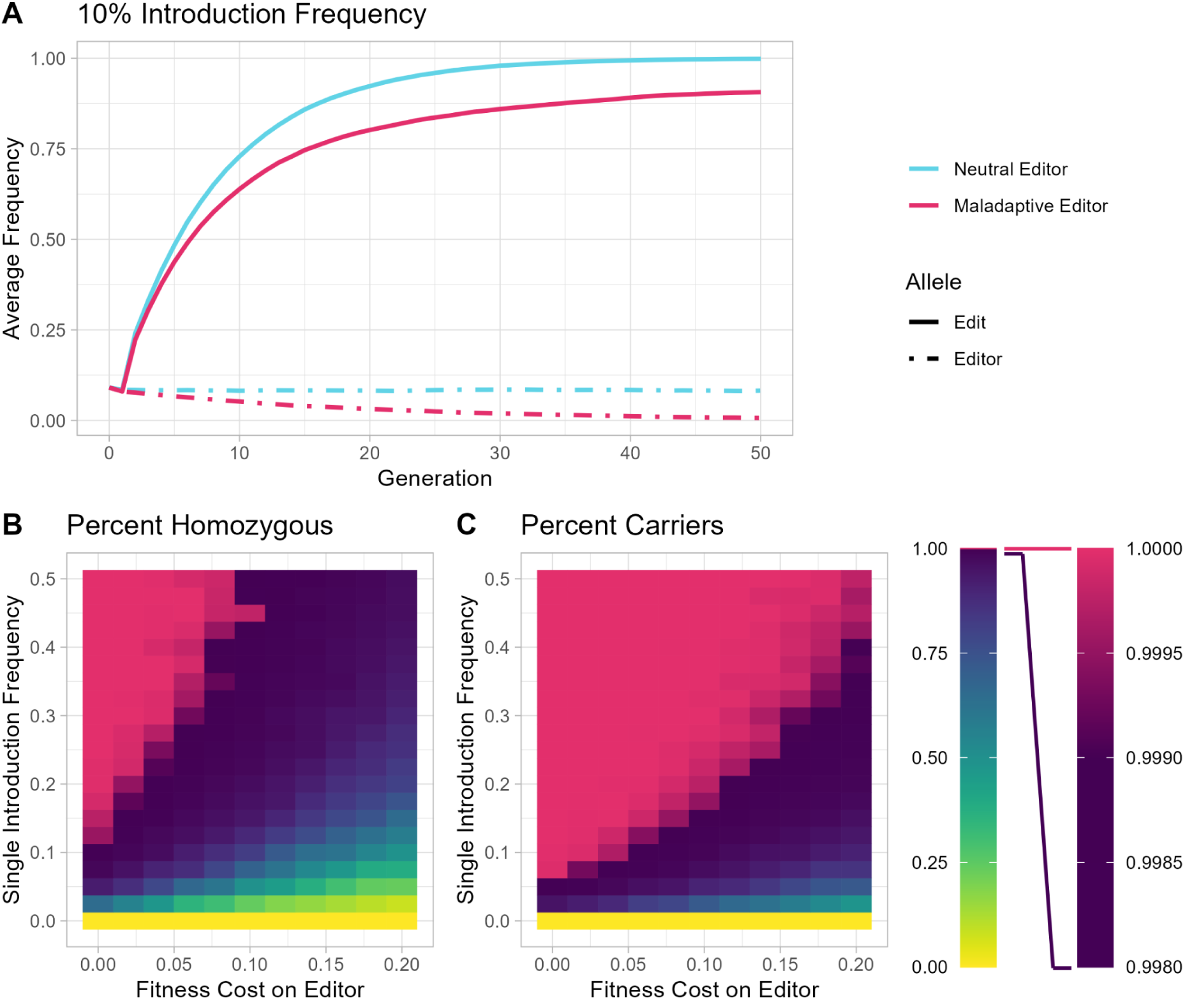
Behavior of a Non-neutral editor & a Neutral edit. **A)** The average allele frequency of edits after being introduced at 10% frequency. Introduced individuals are homozygous for both edit and editor. The non-neutral editor includes an additive 5% fitness cost. **B)** Average percent of the population that is homozygous for our edit after 50 generations, for various editor introduction frequencies and fitness costs associated with the editor. **C)** Average percent of the population that carries at least one copy of the edit, for various editor introduction frequencies and fitness costs.

**Supplementary Figure 4.**
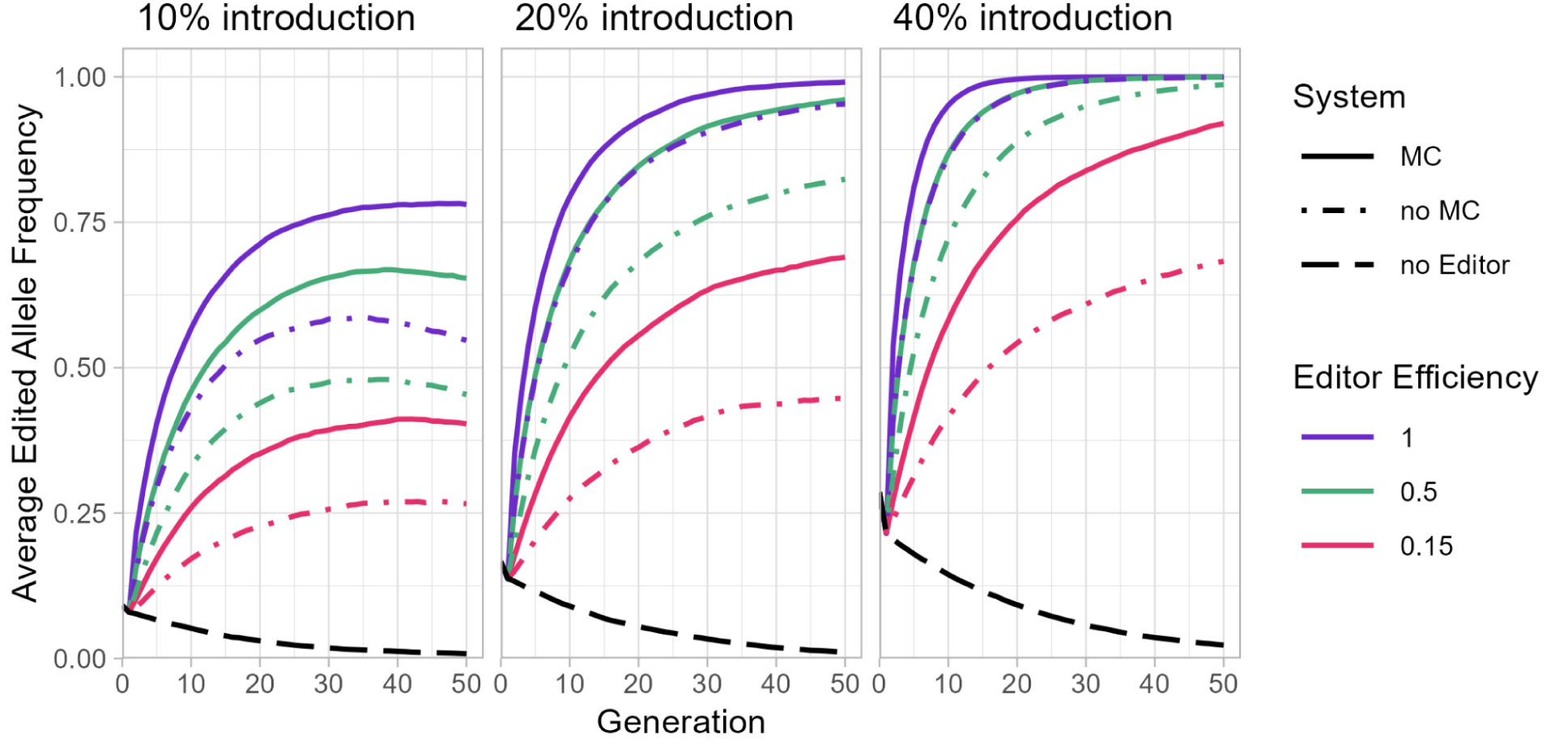
Increasing Introduction Frequency can Overcome Fitness Costs when Editor Efficiency is low. The average edited allele frequency over time when each copy of the edited allele confers a 5% fitness cost. Releases consist of all males homozygous for both the editor and the edit (or only the edit, in the case of the no Editor system). The systems with editor introduced either have maternal carryover (MC) or do not have maternal carryover occurring (no MC).

**Supplementary Figure 5.**
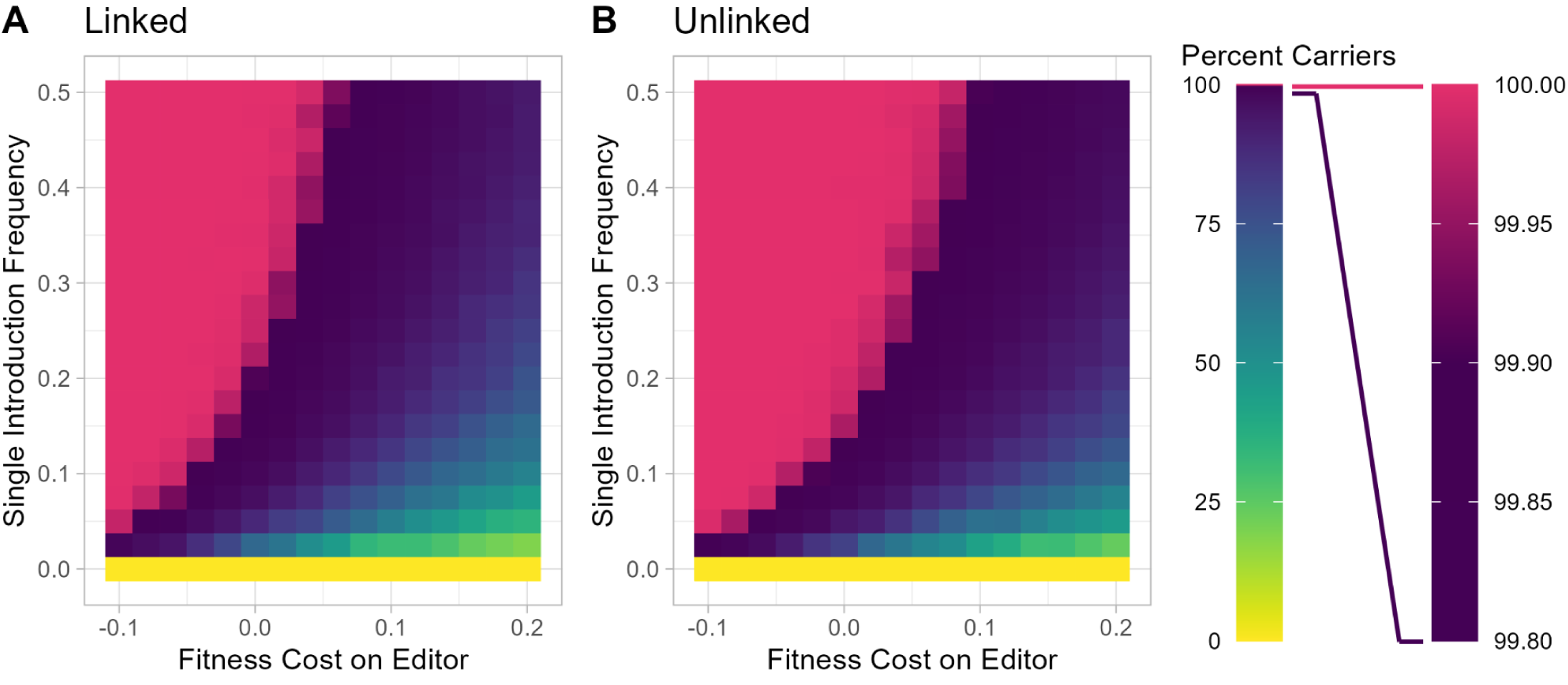
Effects of Linkage on Population Modification, with Cost on the Editor. Rates of germline editing are 50% and there is no maternal carryover editing. No costs are associated with the edit. **A, B)** The percentage of the population that has at least one edited allele / sail present in their genome, averaged over 20 simulations. For moderate costs, linkage between editor and edits results in lower rates of editing **(A)** than does independent assortment. **(B)**.

**Supplementary Figure 6.**
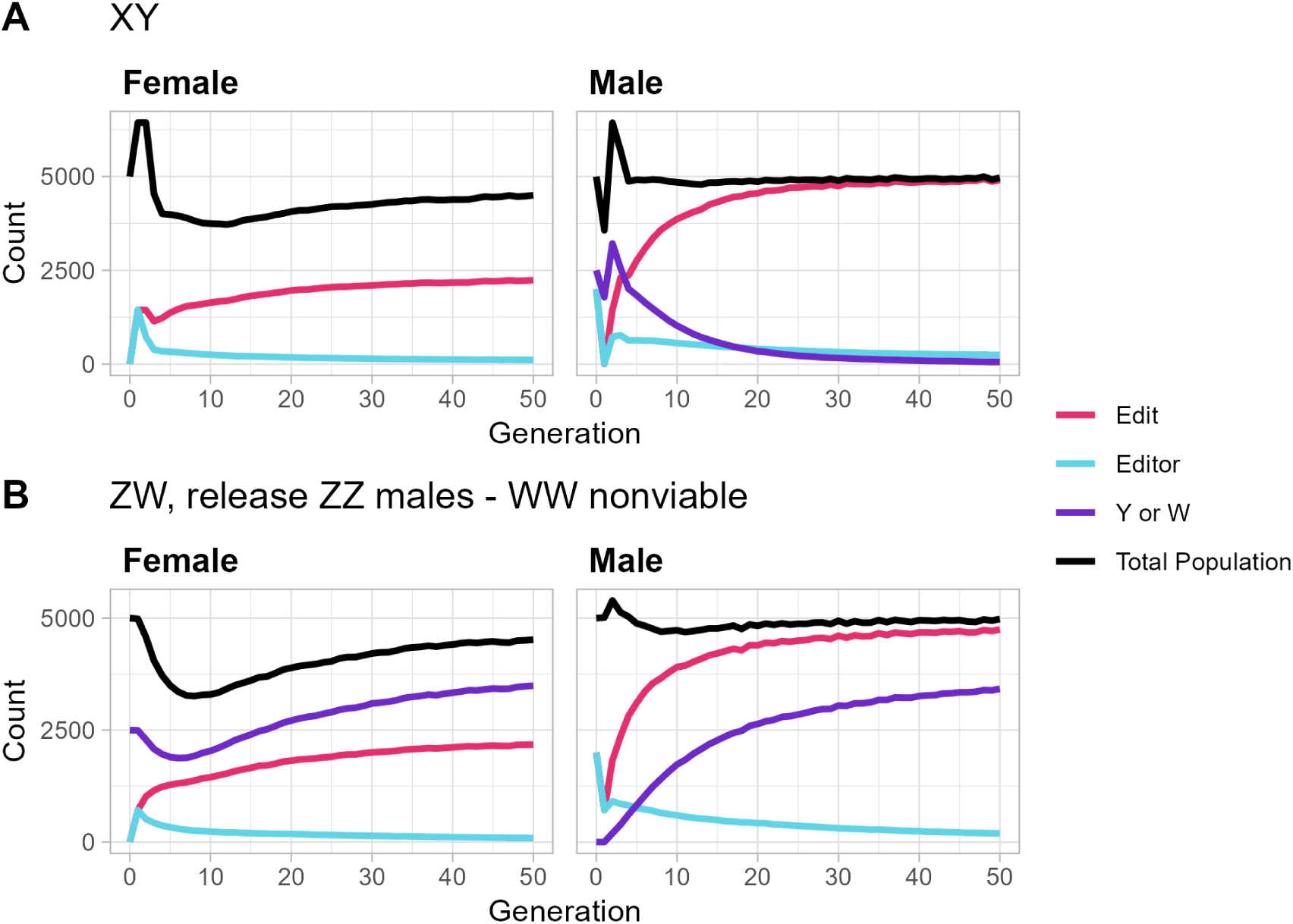
Amounts of Edit and Editor by Sex. **A)** Average total population graphed with the average number of edit-bearing, editor-bearing, and Y-bearing individuals. The editor is introduced in males, is present in only females for the first generation, and then drops quickly in females. The low level of editor in females is because editor offspring are skewed male. The higher frequency of the editor in males over females results in the editor being lost over time, because the editor is removed due to dilution: as sex distortion increases, the editor is segregated into males and fewer of these–as compared with non-editor males– are chosen to participate in mating with the Non-editor females. **B)** The same as **A**, for a ZW system where WW is nonviable. As with the XY system in **A**, the editor has a higher frequency in males than females and decreases over time.

**Supplementary Figure 7:**
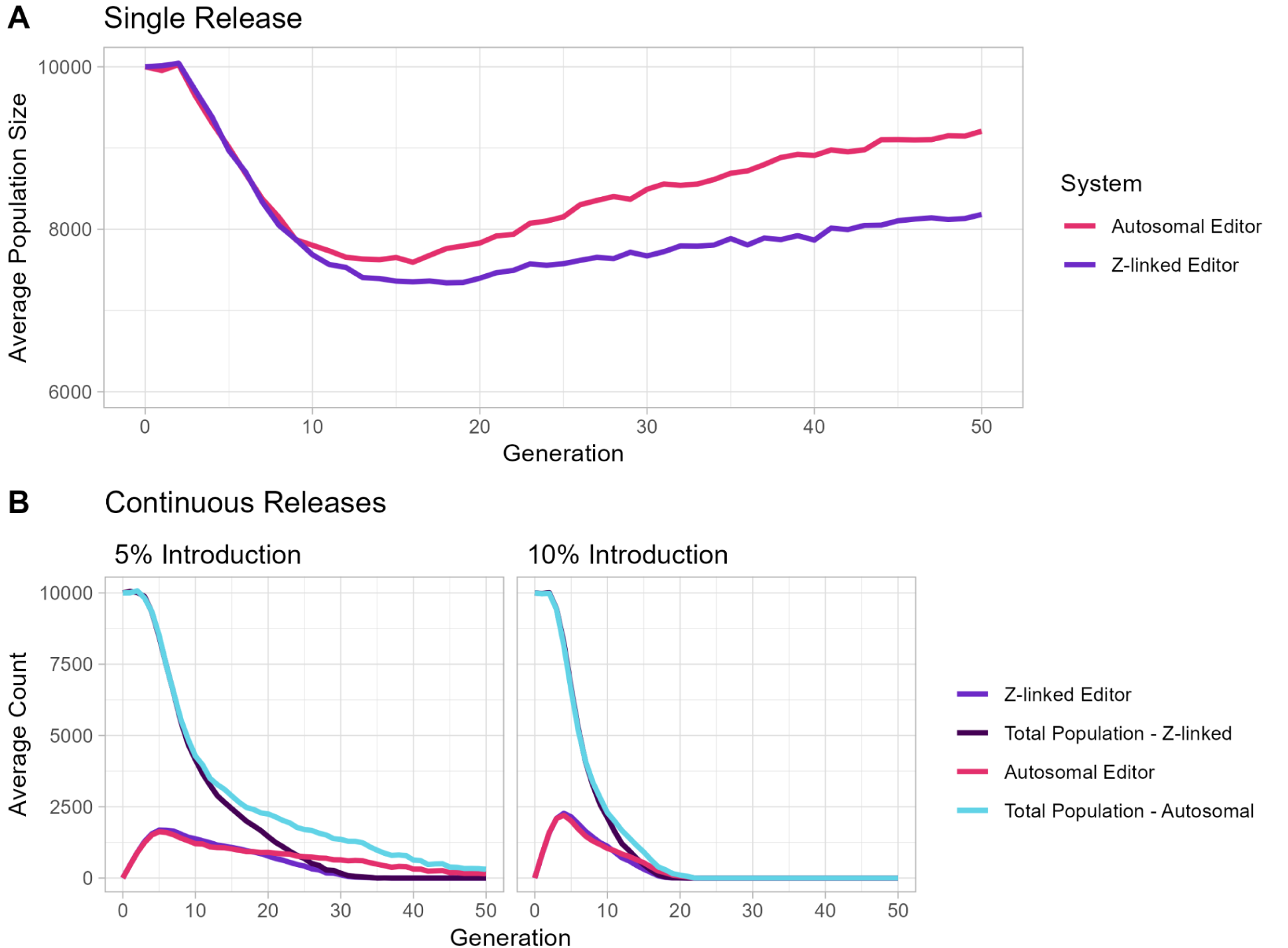
Suppression Behavior of a Z-linked Aromatase Editor. **A)** The average population size over time, following a single release of editor-bearing individuals at 10% of the carrying capacity. Since the released individuals are males, and population numbers are determined by the number of fertile females, these additional males are not counted at generation 1. **B)** The average population size and the number of editor alleles over time. Individuals are released every generation at either 5% or 10% of the carrying capacity. Those released individuals do not count towards the number of alleles in the population, but all of their offspring do.

**Supplementary Figure 8.**
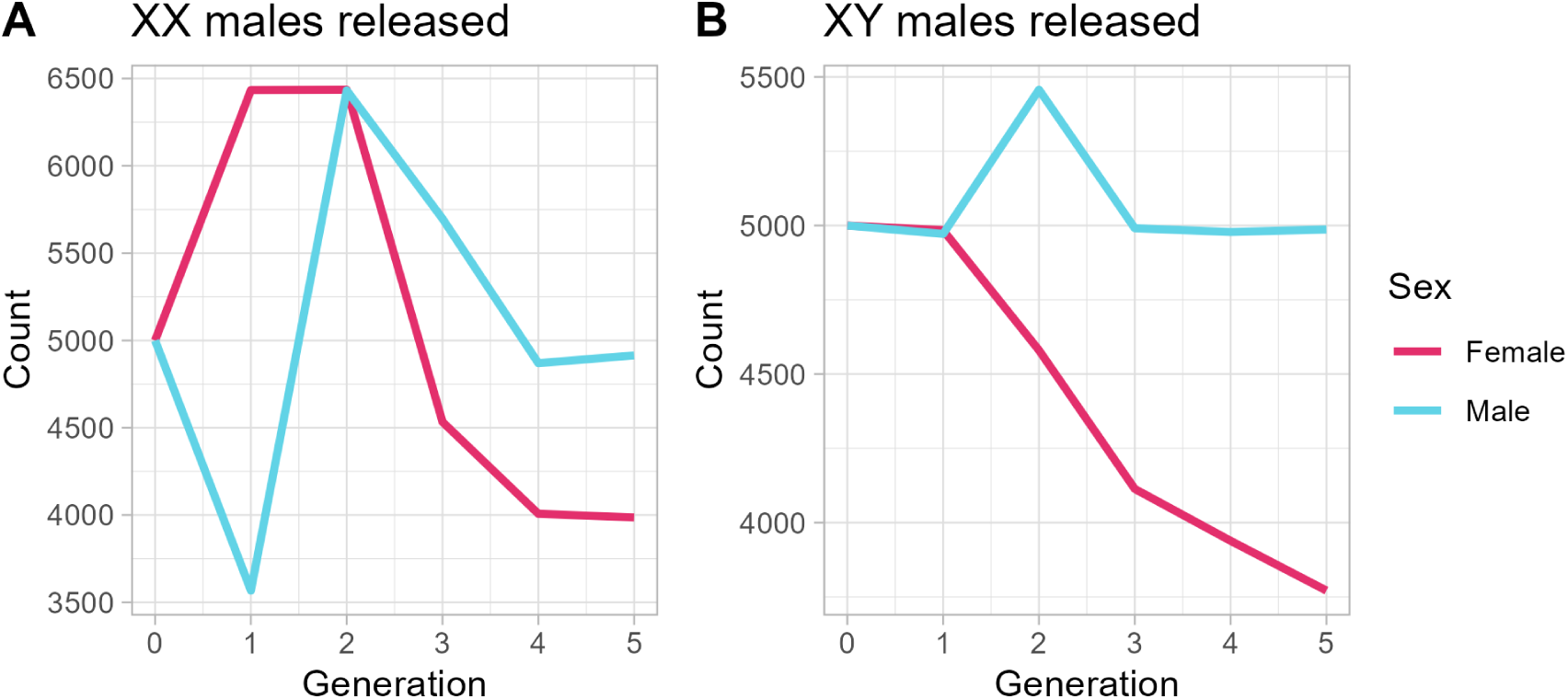
Amount of Males and Females under Population Suppression. **A)** XX males carrying both editor and edit are released into the population at a frequency of 20%, at Generation 0. Notice that after this release, the number of females initially spikes due to increased X frequency. In the second generation, the number of males and females are equal, and higher than carrying capacity due to an increased number of females. After this, the frequency of the edit increases in the population and the sex ratio becomes skewed towards male. **B)** Under the same conditions as A, XY males are released. Notice that there is almost immediate sex skew and no spike in population, because there is no change in the frequency of X and Y chromosomes.

**Supplementary Figure 9:**
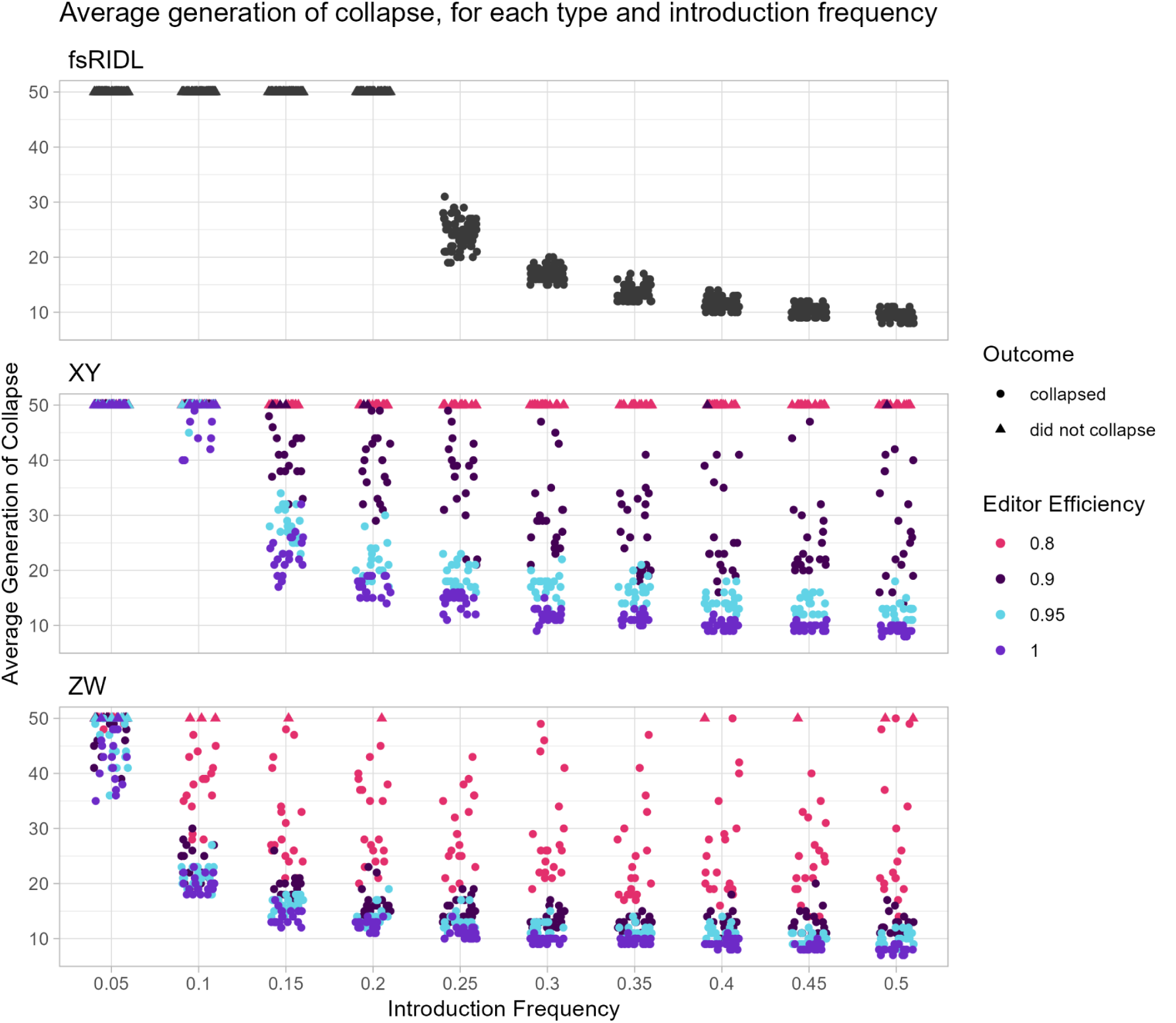
Average time to Population Collapse, for various editor efficiencies. Each dot represents the last non-zero generation of a simulation using the given editor efficiency. As such, simulations that ran to generation 50 but did not collapse are also shown here, but are represented by triangles instead of circles. Twenty simulations were run for each scenario, i.e., for each editor efficiency and system. In the fsRIDL case, changing the editor efficiency did not affect the simulation, as there is no editor present. Even so, there are 80 points for each introduction frequency.

**Supplementary Figure 10.**
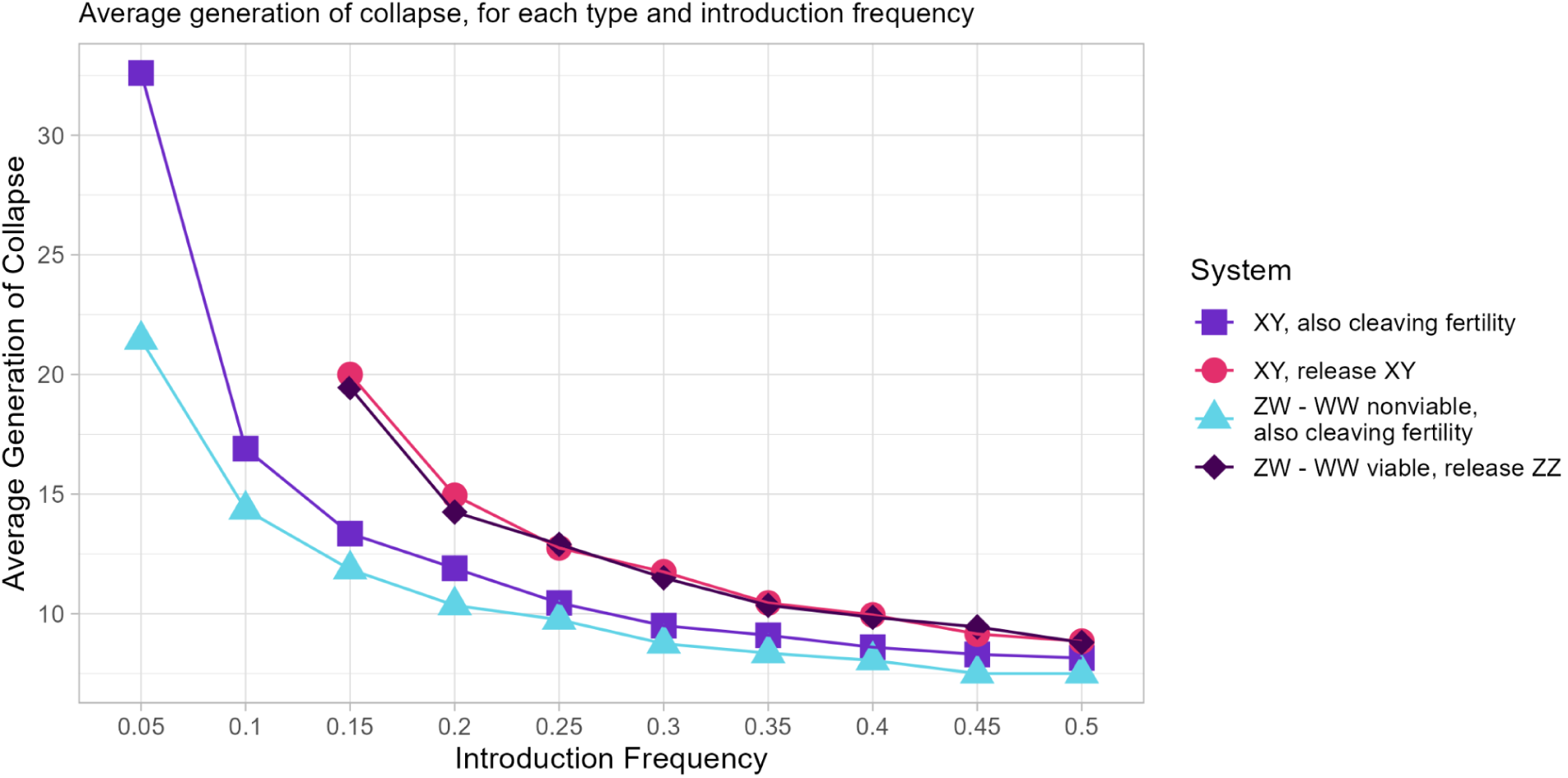
Average time to Population Collapse, including addition of an edit that sterilizes females when homozygous. The same as Figure 6B from the main text, but for different sex determination systems and varying modifications. Points plotted here are the average of 20 simulations, where all 20 simulations went to collapse within 50 generations. If all 20 simulations did not go to fixation within 50 generations, the corresponding point is not plotted here. Transgenic individuals were introduced at the corresponding introduction frequency, once every generation, until collapse. Purple squares (XY system) and light blue triangles (ZW system) show that germine cleavage of a haplosufficient gene required in somatic cells for female fertility, as well as a gene required for femalness, significantly increases the efficiency of an Allele Sail as a suppressive mechanism.

